# Determination of the factors responsible for host tropism of SARS-CoV-2-related bat coronaviruses

**DOI:** 10.1101/2023.04.13.536832

**Authors:** Shigeru Fujita, Yusuke Kosugi, Izumi Kimura, Kenzo Tokunaga, Jumpei Ito, Kei Sato

**Author notes:** Contributed equally to this study (Shigeru Fujita, Yusuke Kosugi). Corresponding author (Kei Sato). **Conflict of interest:** The authors declare that no competing interests exist.

## Abstract

Differences in host ACE2 genes may affect the host range of SARS-CoV-2-related coronaviruses (SC2r-CoVs) and further determine the tropism of host ACE2 for the infection receptor. However, the factor(s) responsible for determining the host tropism of SC2r-CoVs, which may in part be determined by the tropism of host ACE2 usage, remains unclear. Here, we use the pseudoviruses with the spike proteins of two Laotian SC2r-CoVs, BANAL-20-236 and BANAL-20-52, and the cells expressing ACE2 proteins of eight different Rhinolophus bat species, and show that these two spikes have different tropisms for Rhinolophus bat ACE2. Through structural analysis and cell culture experiments, we demonstrate that this tropism is determined by residue 493 of the spike and residues 31 and 35 of ACE2. Our results suggest that SC2r-CoVs exhibit differential ACE2 tropism, which may be driven by adaptation to different Rhinolophus bat ACE2 proteins.

## Introduction

A series of SARS-CoV-2-related coronaviruses (SC2r-CoVs), which are phylogenetically related to SARS-CoV-2, were identified in Rhinolophus bats in China and Southeast Asian countries. For example, RaTG13 (*Rhinolophus affinis* in China, 2013)^1^, RmYN02 (*R. malayanus* in China, 2019)^2^, RacCS203 (*R. acuminatus* in Thailand)^3^, RpYN06 (*R. pusillus* in China, 2020)^4^, RshSTT182 (*R. shameli* in Cambodia, 2010)^5^, BANAL-20-236 (B236; *R. marshalli* in Laos, 2020)^6^, and Rc-o319 (*R. cornutus* in 2013, Japan)^7^ were reported so far. Also, SC2r-CoVs, such as Pangolin-CoV and MpCoV-GX were identified in pangolins^8, 9^. These findings support the concept that SARS-CoV-2 emergence is caused by the spillover of certain SC2r-CoV into humans. Importantly, the spike (S) proteins of some SC2r-CoVs, such as RaTG13 (ref.^10^), B236 (ref.^6^) and those identified in pangolins^11, 12^ are capable of binding to human angiotensin converting enzyme 2 (ACE2), the receptor for SARS-CoV-2 infection. Therefore, it is plausible to assume that some SC2r-CoVs, which can bind to human ACE2, circulate in Rhinolophus bats and pangolins in the wild, particularly those residing in Southeast Asian countries. However, because bat *ACE2* genes are highly diversified^13–15^, it is hypothesized that the difference of bat *ACE2* genes can affect the host range of SC2r-CoVs and further modulate the tropism of host ACE2 for the infection receptor.

B236 is a replication-competent SC2r-CoV that was isolated from rectal swabs of Laotian *R. marshalli* by Temmam et al.^6^ In this previous study, the viral sequences of BANAL-20-52 (B52) and BANAL-20-103 (B103) were identified from the samples of *R. malayanus* and *R. pusillus*^6^. Importantly, these three BANAL-20 viruses are phylogenetically close to SARS-CoV-2 (ref.^6^). The amino acid sequences of S receptor binding motif of B52 and B103 are identical, and the receptor binding domain (RBD) of B52/103 RBD more strongly binds to human ACE2 than SARS-CoV-2 S RBD^6^. These observations suggest that B236, B52 and B103 are capable of using human ACE2 for the infection receptor, however, the host tropism of these viruses, which can be determined in part by the tropism of host ACE2 usage, remains unclear. In this study, we particularly focus on the two Laotian SC2r-CoVs, B236 and B52, and elucidate the difference of host ACE2 tropism.

## Results

### Difference of ACE2 tropism between B236 and B52

We set out to understand the phylogenetic relationship of SC2r-CoVs in Rhinolophus bats and pangolins. Consistent with a previous report^6^, most of the SC2r-CoVs that are capable of using human ACE2 for the infection receptor, formed a cluster with SARS-CoV-2 (**Fig. 1A**).

**Fig. 1.**
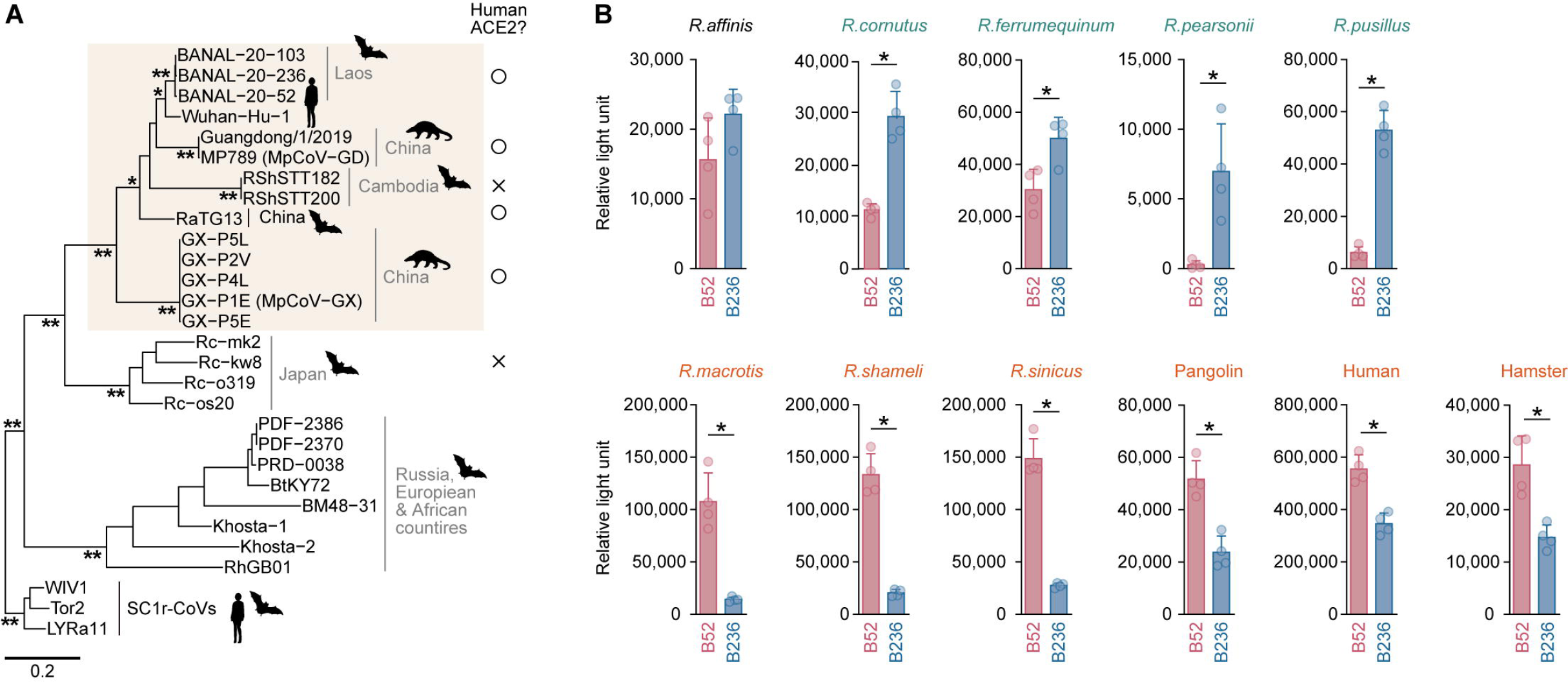
Different ACE2 tropism of the two Laotian SC2r-CoVs. **(A)** Maximum likelihood tree of SC2r-CoVs and SARS-CoV-2 (strain Wuhan-Hu-1) based on their nucleotide sequences corresponding to RBD in S. SARS-CoV-1 (strain Tor2) and two SARS-CoV-1-related coronaviruses (SC1r-CoVs; WIV1 and LYRa11) are included as an outgroup. *, >0.8 bootstrap value; **, >0.9 bootstrap value. Scale bar indicates genetic distance. The usability of human ACE2 for SC2r-CoVs infection is indicated with ○ (yes) or × (no), respectively, and the clade of SC2r-CoVs that can use human ACE2 is shaded in brown. **(B)** Pseudovirus assay. HIV-1-based reporter viruses pseudotyped with the S proteins of B52 or B236 were prepared. The pseudoviruses were inoculated into a series of HOS-TMPRSS2 cells stably expressing Rhinolophus bat ACE2 cells at 1 ng HIV-1 p24 antigen. The infectivity (relative light unit) in each target cell is shown. The host species in which ACE2 is preferred by B52 or B236 are indicated in green and orange, respectively. Data are expressed as the mean with SD. Assays were performed in quadruplicate. Statistically significant differences (\**P* < 0.05) between B52 or B236 were determined by two-sided Student’s t test.

To investigate the host ACE2 tropism of two Laotian SC2r-CoVs, B236 and B52, we prepared the HOS-TMPRSS2 cell lines that stably express ACE2 proteins of eight Rhinolophus bat species: *R. affinis, R. cornutus, R. ferrumequinum, R. macrotis, R. pearsonii, R. pusillus, R. shameli,* and *R. sinicus*, most of which are found throughout Southeast and East Asia (**Fig. S1**). As controls, we also prepared the HOS-TMPRSS2 cells stably expressing ACE2 proteins of human, hamster and pangolin. We then prepared the human immunodeficiency virus type 1 (HIV-1)-based pseudoviruses with the S proteins of B236 and B52 and inoculated them into a series of target cells prepared. As shown in **Fig. 1B**, the HOS-TMPRSS2 cells expressing *R. affinis* ACE2 exhibited similar infectivity to both B236 and B52. On the other hand, B236 exhibited higher infectivity than B52 in the cells expressing ACE2 proteins of *R. cornutus, R. ferrumequinum, R. pearsonii, R. pusillus* (**Fig. 1B**). In contrast, B52 exhibited higher infectivity than B236 in the cells expressing ACE2 proteins of *R. macrotis, R. shameli, R. sinicus* as well as those expressing ACE2 proteins of pangolin, human and hamster (**Fig. 1B**). These results suggest that the ACE2 tropism of B236 and B52 is different among animal species.

### Interspecies polymorphisms of ACE2 linked to the susceptibility to B52 than B236 infection

We next investigated the determinant factor(s) that are responsible for the ACE2 tropism of B52 and B236 on both sides of hosts (i.e., ACE2) and viruses (i.e., viral S). We first addressed the host side and assumed that the evolutionary relationship of horseshoe bat species. However, there is no clear correlation between the tropism of B236/B52 and the phylogenetic relationship of the host (**Fig. S2A**). Also, the phylogenetic tree of ACE2 gene did not show a clear association with the tropism of B236/B52 (**Fig. S2B**). For example, *R. macrotis* ACE2 gene is phylogenetically closely related to *R. pusillus* and *R. cornutus* (**Fig. S2B**). However, *R. macrotis* ACE2 showed higher susceptibility to B52 than B236, while the ACE2 proteins of *R. pusillus* and *R. cornutus* displayed higher susceptibility to B236 than B52 (**Fig. 1B**). These observations suggest that the differences in susceptibility of ACE2 proteins to B52 and B236 could not solely be explained by the phylogenetic relationships of the host species and ACE2 gene (**Fig. S2**).

To identify the genetic determinants of susceptibility to B52 and B236 infections in ACE2 proteins, we then assessed the amino acid polymorphisms in ACE2 proteins that can be associated with susceptibility to B52 and B236. As shown in **Fig. 2A**, the susceptibility of ACE2 proteins to B236 exhibited a strong inverse correlation with that of B52 (except for the ACE2 proteins of human and *R. pearsonii*). We calculated the relative infectivity score between B236 and B52 [i.e., log_10_(B236 infectivity/B52 infectivity)] for each ACE2, and subsequently, evaluated the association between this score and amino acid polymorphism at each site in ACE2 protein. For the analysis, the data for outlier species, human and *R. pearsonii*, were excluded. At the permissive statical threshold (*P* < 0.1), we detected eight amino acid sites associated with the susceptibility (**Fig. 2B and 2C**). Of these, residue 35 exhibited the strongest association (*P* = 0.0067). In these eight residues, four residues positioned at 30, 31, 35 and 79 are located in the region that interacts with SARS-CoV-2 S RBD^16^. Particularly, Shang et al. showed that the residues positioned at 31 (K31) and 35 (E35) of human ACE2 are crucial to form hydrogen bonds with Q493 of SARS-CoV-2 S (**Fig. 2D**)^16^. When we focus on the virus side, the amino acid similarity of the RBDs of B52 and B236 are very high (221/223; 99.1%) and the two amino acid residues positioned at 324 (D324 for B236, E324 for B52) and 493 (K493 for B236, Q493 for B52) are different in the RBD between B52 and B236 (**Fig. S3**). Importantly, the residue 493 is the only residue of SARS-CoV-2 S protein that can interact with human ACE2 and is different between B52 and B236 (**Fig. 2D and S3**). Moreover, the co-crystal structure of B236 S RBD and human ACE2 showed that the K493 of B236 S interacts with the E35 of human ACE2 (ref.^6^). Because a previous paper^16^ and our analyses (**Fig. 2A-2C**) suggested the importance of the residues positioned at 31 and 35 of ACE2 to interact with viral S protein, we addressed the possibility that the residues 31 and 35 of Rhinolophus bat ACE2 interact with the residue 493 of B52/236 S. We prepared homology models of the ACE2 proteins of four Rhinolophus bats, *R.cornutus, R. macrotis, R. pusillus* and *R. sinicus,* and constructed docking models with the S RBDs of B236 and B52. As summarized in **Fig. 2C**, the ACE2 tropism by B236/52 is closely correlated to the two residues positioned at 31 and 35. In fact, the docking models of S RBD and ACE2 showed that the K493 of B236 formed salt bridges with the D31 and E35 of the ACE2 proteins of *R. cornutus* and *R. pusillus*, while the electrostatic repulsion was observed between the K493 of B236 and the K31 and K35 of the ACE2 proteins of *R. macrotis,* and *R. sinicus* (**Fig. 2E**). In contrast, the Q493 of B52 formed hydrogen bonds with the K31 of the ACE2 proteins of *R. macrotis,* and *R. sinicus*, or with D31 of the ACE2 proteins of *R. cornutus* and *R. pusillus* (**Fig. 2F**). These observations suggest that the electrostatic interaction between the residue 493 of B236/B52 S protein and the residues 31 and 35 of host ACE2 protein is associated with the tropism of 52 and B236.

**Fig. 2.**
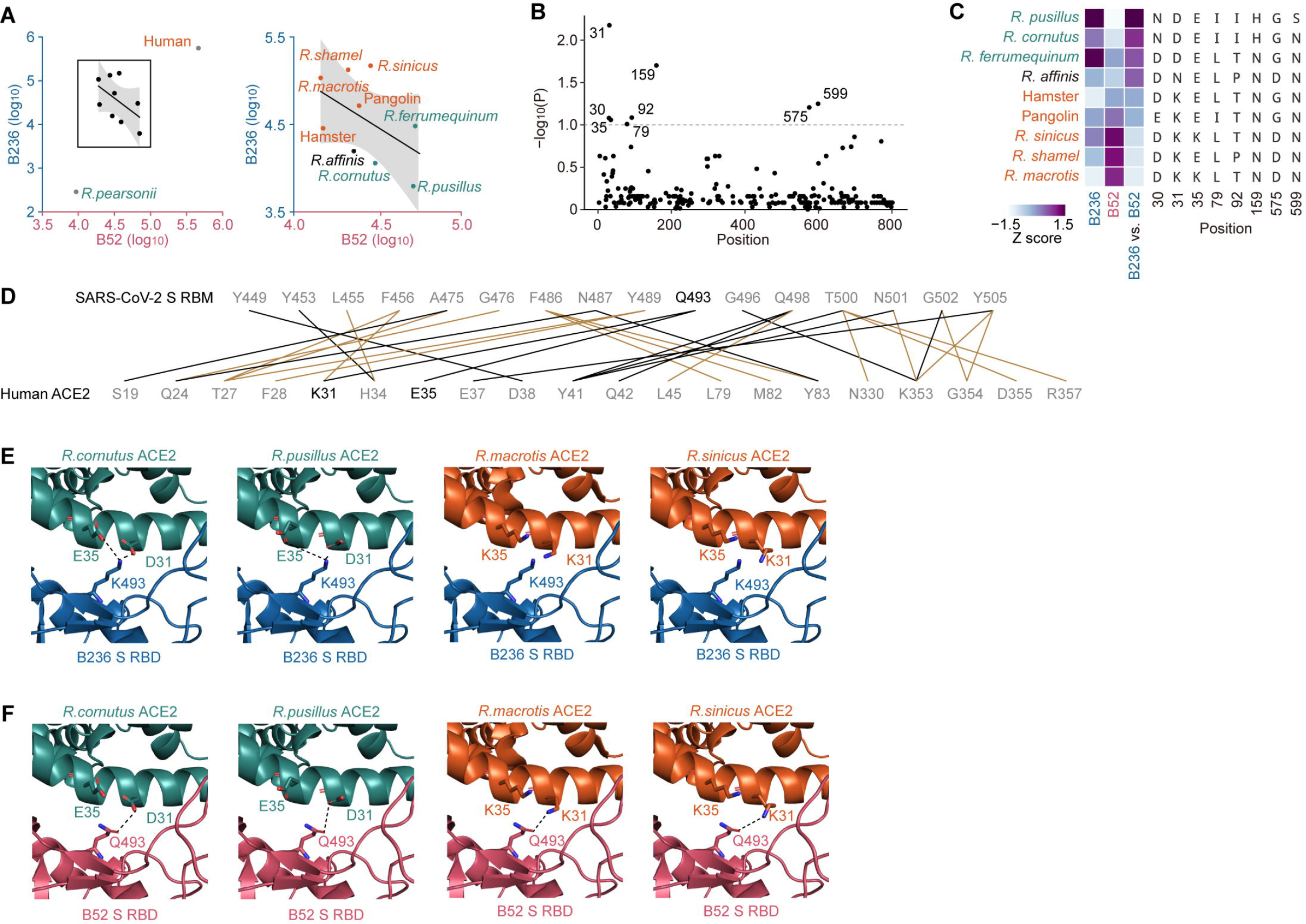
Interaction between Rhinolophus bat ACE2 and the S proteins of two Laotian SC2r-CoVs. **(A)** Inverse correlation of the ACE2 susceptibility to B236 and B52 infection. Boxed region is zoomed in the right panel. **(B)** Association between the B52 and B236 infectivity and ACE2 polymorphism among animal species. The association between the relative infectivity [log_10_(B236 infectivity / B52 infectivity)] for each ACE2 protein and each polymorphic amino acid site was evaluated by one-way ANOVA. Dashed line, *P* = 0.1. Outlier species (human and *R. pearsonii*; gray dots in Fig. 1B) were excluded from the analysis. **(C)** Amino acid sites associated with the B236 and B52 infection tropism. Heatmaps of the Z scores of B236 infectivity, B52 infectivity, and the relative infectivity are shown on the left. (**D–F**) Structural insights into the binding of S RBD and ACE2 proteins. (D) The scheme of interaction between SARS-CoV-2 S receptor binding motif (top) and human ACE2 (bottom). Salt bridge or hydrogen bond is indicated in black, and van der Waals interaction is indicated in brown. Q493 of SARS-CoV-2 S and K31 and E35 of human ACE2 are indicated in black. This information is referred from a previous report^16^. RBM, receptor binding motif. **(E)** The structural model of complex of B236 S RBD (blue) and the homology models ACE2 of *R. cornutus* (leftmost, green), *R. pusillus* (the second from the left, green), *R. macrotis* (the second from the right, orange) or *R. sinicus* (rightmost, orange), respectively. The residue 493 of B236 S RBD and the residues 31 and 35 of ACE2s are indicated as stick model. Dashed lines indicate salt bridges. **(F)** The structural model of complex of B52 S RBD (red) and the homology models ACE2 of *R. cornutus* (leftmost, green), *R. pusillus* (the second from the left, green), *R. macrotis* (the second from the right, orange) or *R. sinicus* (rightmost, orange), respectively. The residue 493 of B52 S RBD and the residues 31 and 35 of ACE2s are indicated as stick model. Dashed lines indicate hydrogen bonds.

### Determination of the different tropism of B236 and B52

To address the possibility that the interaction between the residue 493 of viral S protein and the residues 31 and 35 of host ACE2 protein explains the different tropism of B236 and B52, we prepared the two S derivatives of B236 and B52, which harbor the mutations at residue 493: B236 S K493Q and B52 S Q493K. The mutations at the residue 493 of S protein did not affect to the levels of S proteins incorporated into the released pseudoviral particles (**Fig. 3A**). We then selected the two Rhinolophus ACE2 proteins as representatives, those from *R. pusillus* and *R. macrotis*, which are strongly preferred by B236 and B52, respectively (**Fig. 1B**), for pseudovirus infection experiments. As shown in **Fig. 3B**, in the cells expressing *R. pusillus* ACE2, which is preferred by B236, the infectivity of B236 S pseudovirus was significantly decreased (89.7-fold) by the K493Q substitution. In contrast, the infectivity of B52 S pseudovirus was significantly increased (24.1-fold) by the Q493K substitution (**Fig. 3B**). In the cells expressing *R. macrotis* ACE2, which is preferred by B52, the infectivity of B52 pseudovirus was significantly decreased (2.4-fold) by the Q493K substitution, while that of B236 was 5.9-fold increased by the K493Q mutation (**Fig. 3B**). These results suggest that the residue 493 of B52/236 S determines the tropism to Rhinolophus ACE2.

**Fig. 3.**
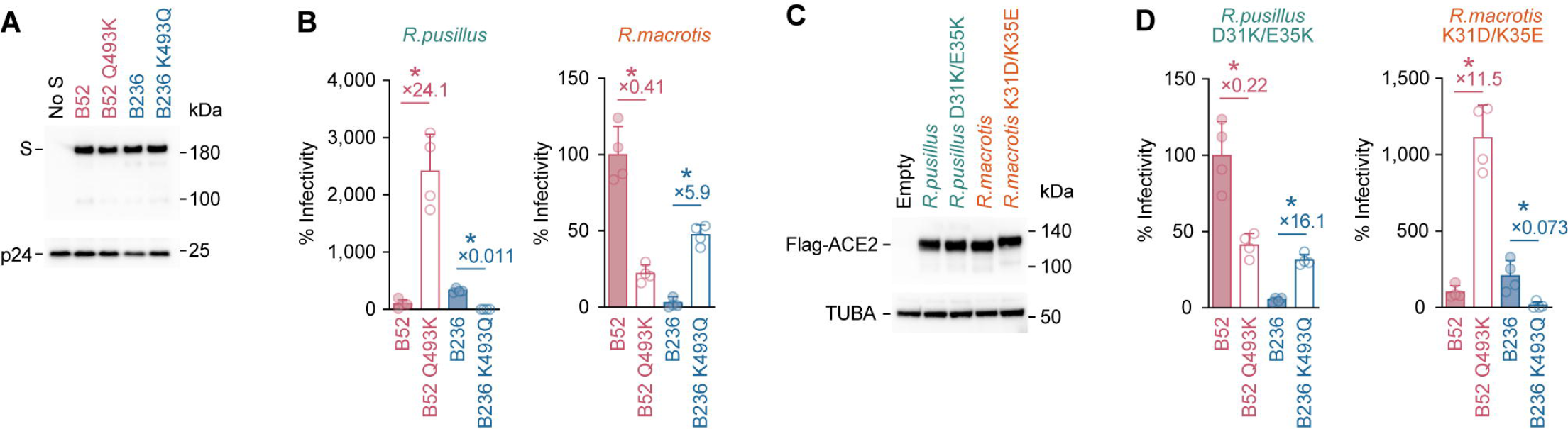
ACE2 tropism determined by the interaction between the residues 31/35 of ACE2 and the residue 493 of S. **(A)** Western blotting. A representative blot of pseudovirus is shown. HIV-1 p24 is an internal control for the pseudovirus. kDa, kilodalton. (**B and D**) Pseudovirus assay. HIV-1-based reporter viruses pseudotyped with the S proteins of B236, B52 or their derivatives were prepared. The pseudoviruses were inoculated into a series of HEK293 cells transiently expressing Rhinolophus bat ACE2 cells at 2 ng HIV-1 p24 antigen, and the percentages of infectivity compared to that of the virus pseudotyped with B52 are shown. The number in the panel indicate the fold change of the B236 value to the B52 value in each target cell. (**C**) Western blotting. A representative blot of the cells transiently expressing flag-tagged Rhinolophus bat ACE2 cells are shown. TUBA is an internal control for the cells. kDa, kilodalton. In (**B**) and (**D**), data are expressed as the mean with SD. Assays were performed in quadruplicate. The number in the panel indicate the fold change versus parental S. Statistically significant differences (\**P* < 0.05) between B236 and B52 were determined by two-sided Student’s t test.

To further address the possibility that the residues 31 and 35 of ACE2 receptor are responsible for the tropism of B236/52, we generated the plasmids expressing *R. pusillus* ACE2 D31K/E35K and *R. macrotis* ACE2 K31D/K35E. The mutations at residues 31 and 35 of Rhinolophus ACE2 did not affect to their protein expression levels (**Fig. 3C**). In the case of *R. pusillus* ACE2 D31K/E35K, the B52 infectivity was 4.5-fold decreased by the Q493K substitution, while the B236 infectivity was 16.1-fold increased by the K493Q substitution (**Fig. 3D**). In the case of *R. macrotis* ACE2 K31D/K35E, the B52 infectivity was 11.5-fold increased by the Q493K substitution, while the B236 infectivity was 13.7-fold decreased by the K493Q substitution (**Fig. 3D**). Altogether, these findings suggest that the ACE2 tropism of B236 and B52 is determined by the residue 493 of viral S proteins and the residues 31 and 35 of ACE2 receptors are responsible for the viral tropism.

## Discussion

In this study, we showed that the SC2r-CoVs identified in Laotian bats, B236 and B52, exhibit different ACE2 tropism. We further demonstrate that this tropism difference is determined by the amino acid residue positioned at 493 of their S proteins. Structural analysis suggests that residue 493 of the viral S protein plays a critical role in the interaction mediated by salt bridge with amino acid residues positioned at 31 and 35 of the host ACE2 protein. Our results provide insight into the host tropism of SC2r-CoVs, which is defined by the ACE2 receptor.

Interactions between viral proteins and host receptors that determine host range are known from other viruses (reviewed in ref.^17^). For example, the host tropism of influenza A viruses (IAVs) is determined by the affinity of the viral haemagglutinin for the sialic acids of the host species: human IAVs prefer to bind to α2-3-linked sialic acid, whereas avian IAVs prefer to bind to α2-6-linked sialic acid. (reviewed in ref.^18, 19^). In primate lentiviruses (PLVs), including human immunodeficiency virus type 1 (HIV-1), the viral envelope protein (Env) binds to two host receptor proteins, the CD4 protein (major receptor) and chemokine receptors (coreceptors), to initiate infection. Russel et al. showed that CD4 receptor diversity is an ancient protective mechanism against PLVs^20^. Moreover, although HIV-1 and related primate lentiviruses use CCR5 as the infection coreceptor, some PLVs that are evolutionarily unrelated to HIV-1 use CCR2 (ref.^21, 22^) or CXCR6 (ref.^23–25^). Furthermore, the host range of the Ebola virus is determined by the difference in amino acid residues in the Niemann-Pick C1 protein, the infection receptor^26, 27^. Therefore, the identification of the amino acid residues of the viral S protein that determine the specificity of the host receptor proteins leads to an estimate of the host range of the virus. Further, if the use of human ACE2 and the amino acid residues of the SC2r-CoV S proteins that determine its affinity can be identified, it will be possible to infer from the *S* gene sequence alone whether the SC2r-CoV of concern is capable of spreading to humans.

The mutation pattern of the residue 493 of SARS-CoV-2 S is extraordinary: the S proteins of prior SARS-CoV-2 variants of concern (VOCs) including ancestral Wuhan-Hu-1 strain harbor Q493 (ref.^28^), while those of the two major SARS-CoV-2 VOCs, Omicron BA.1 and BA.2, harbor R493 (ref.^29^). Interestingly, the R493 has reverted to Q493 in subsequent Omicron subvariants such as Omicron BA.4, BA.5, BA.2.75, BQ.1.1 (ref.^30–33^). Regarding this, Wang et al. showed that the R493Q reversion mutation improves the affinity to human ACE2 (ref.^34^). Additionally, we have previously shown that the R493Q mutation contributes to the evasion of humoral immunity induced by BA.2, which carries R493, in rodents^30^. Therefore, residue 493 of SC2r-CoVs may be relatively easily converted to R493 and Q493, depending on immune evasion and/or affinity to the host ACE2 protein, and the humoral immunity of Rhinolophus bats against SC2r-CoVs may be associated with the conversion of residue 493.

### Limitation of this study

Here we focused on two Laotian SC2r-CoVs, B236 and B52. However, as the *ACE2* genes of their host Rhinolophus bats, *R. marshalli* and *R. malayanus*, have not yet been determined, we could not test the receptor tropism of these two SC2r-CoVs on their host species. As there are still several Rhinolophus bat species in Southeast Asia, it would be possible to map the range and diversity of each SC2r-CoV by studying its habitat and ACE2 gene diversity.

In conclusion, here we showed that *ACE2* genes are diversified in Rhinolophus bats and this may determine the tropism of SC2r-CoVs to Rhinolophus bat species. The difference of SC2r-CoVs in Rhinolophus ACE2 tropism may be a driving force that promotes the diversity of circulating viruses in Rhinolophus bats and further confers infectivity to a variety of host species, including humans. The previous studies focusing on the SARS-CoV-2 VOCs have shown that the host range can be altered by mutations in the *S* gene. For example, although the S protein of the ancestral Wuhan-Hu-1 strain is unable to use murine ACE2 as an infection receptor, the N501Y mutation present in the S proteins of the Alpha and subsequent variants allows murine ACE2 to be used for infection^35^. Identifying mutations in the S protein that determine host receptor usage should be an important study to estimate the host range of SC2r-CoVs of concern. Further investigations will be needed to identify and predict SC2r-CoVs in the wild that can be transmitted to humans.

## Author Contributions

Jumpei Ito performed phylogenetic and statistical analysis.

Shigeru Fujita, Yusuke Kosugi, Izumi Kimura and Kenzo Tokunaga performed cell culture experiments.

Yusuke Kosugi prepared protein structure models. Jumpei Ito and Kei Sato wrote the original manuscript.

All authors designed the experiments, interpreted the results, and reviewed and proofread the manuscript.

The Genotype to Phenotype Japan (G2P-Japan) Consortium contributed to the project administration.

## Acknowledgments

We would like to thank all members belonging to The Genotype to Phenotype Japan (G2P-Japan) Consortium. The super-computing resource was provided by Human Genome Center at The University of Tokyo. We thank Dr. Shin Murakami (The University of Tokyo) for providing pCAGGS-blast-RcACE2.

This study was supported in part by AMED SCARDA Japan Initiative for World-leading Vaccine Research and Development Centers “UTOPIA” (JP223fa627001, to Kei Sato), AMED SCARDA Program on R&D of new generation vaccine including new modality application (JP223fa727002, to Kei Sato); AMED Research Program on Emerging and Re-emerging Infectious Diseases (JP22fk0108511, to G2P-Japan Consortium and Kei Sato; JP22fk0108516, to Kei Sato; JP22fk0108506, to Kei Sato; JP22fk0108146, to Kei Sato; JP21fk0108494 to G2P-Japan Consortium and Kei Sato; JP21fk0108425, to Kei Sato; JP21fk0108432, to Kei Sato); AMED Research Program on HIV/AIDS (JP22fk0410039, to Kei Sato); JST PRESTO (JPMJPR22R1, to Jumpei Ito); JST CREST (JPMJCR20H4, to Kei Sato); JSPS KAKENHI Grant-in-Aid for Early-Career Scientists (20K15767, to Jumpei Ito; 23K14526, to Jumpei Ito); JSPS Core-to-Core Program (A. Advanced Research Networks) (JPJSCCA20190008, to Kei Sato);); JST SPRING (JPMJSP2108, to Shigeru Fujita).

## Consortia

Keita Matsuno^8^, Naganori Nao^8^, Hirofumi Sawa^8^, Shinya Tanaka^8^, Masumi Tsuda^8^, Lei Wang^8^, Yoshikata Oda^8^, Zannatul Ferdous^8^, Kenji Shishido^8^, Takasuke Fukuhara^8^, Tomokazu Tamura^8^, Rigel Suzuki^8^, Saori Suzuki^8^, Hayato Ito^8^, Yu Kaku^1^, Naoko Misawa^1^, Arnon Plianchaisuk^1^, Ziyi Guo^1^, Alfredo A. Hinay, Jr.^1^, Keiya Uriu^1^, Jarel Elgin M. Tolentino^1^, Luo Chen^1^, Lin Pan^1^, Mai Suganami^1^, Mika Chiba^1^, Ryo Yoshimura^1^, Kyoko Yasuda^1^, Keiko Iida^1^, Naomi Ohsumi^1^, Adam P. Strange^1^, Kazuhisa Yoshimura^9^, Kenji Sadamasu^9^, Mami Nagashima^9^, Hiroyuki Asakura^9^, Isao Yoshida^9^, So Nakagawa^10^, Kotaro Shirakawa^11^, Akifumi Takaori-Kondo^11^, Kayoko Nagata^11^, Ryosuke Nomura^11^, Yoshihito Horisawa^11^, Yusuke Tashiro^11^, Yugo Kawai^11^, Kazuo Takayama^11^, Rina Hashimoto^11^, Sayaka Deguchi^11^, Yukio Watanabe^11^, Ayaka Sakamoto^11^, Naoko Yasuhara^11^, Takao Hashiguchi^11^, Tateki Suzuki^11^, Kanako Kimura^11^, Jiei Sasaki^11^, Yukari Nakajima^11^, Hisano Yajima^11^, Takashi Irie^12^, Ryoko Kawabata^12^, Kaori Tabata^13^, Terumasa Ikeda^13^, Hesham Nasser^13^, Ryo Shimizu^13^, MST Monira Begum^13^, Michael Jonathan^13^, Yuka Mugita^13^, Otowa Takahashi^13^, Kimiko Ichihara^13^, Chihiro Motozono^13^, Takamasa Ueno^13^, Mako Toyoda^13^, Akatsuki Saito^14^, Maya Shofa^14^, Yuki Shibatani^14^, Tomoko Nishiuchi^14^

## Footnotes

8 Hokkaido University, Japan.

9 Tokyo Metropolitan Institute of Public Health, Japan

10 Tokai University, Japan

11 Kyoto University, Japan

12 Hiroshima University, Japan

13 Kyushu University, Japan

13 Kumamoto University, Japan

14 University of Miyazaki, Japan

---

**Table 1.** Summary of the residues 31 and 35 of ACE2 and the tropism of B236 and B52

|  | residue |  |  |
| --- | --- | --- | --- |
| The bat species of ACE2 | 31 | 35 | ACE2 tropism of B236 and B52 |
| *R.cornutus* | D | E | B52 < B236 |
| *R.pusillus* | D | E | B52 < B236 |
| *R.macrotis* | K | K | B52 > B236 |
| *R.sinicus* | K | K | B52 > B236 |
| *R.shameli* | K | E | B52 > B236 |

**Fig. S1. Geological distributions of Rhinolophus bat species.**

The habitat information originates from The IUCN Red List of Threatened Species website (https://www.iucnredlist.org/).

**Fig. S2. Phylogenetic relationship of Rhinolophus bat species.**

**(A)** Time-calibrated species tree for Rhinolophus bat species generated by TimeTree5 (ref.^36^). MYA, million years ago. **(B)** Maximum likelihood tree of Rhinolophus bat ACE2 sequences. Scale bar indicates genetic distance.

**Fig. S3. Amino acid alignment of the RBDs of B52 and B236.**

Residues with nonsynonymous substitution between B52 and B236 are shaded in gray. The alignment was plotted by Multiple Align Show (https://www.bioinformatics.org/sms/index.html).

## Methods

### Nucleotide sequence data collection

The whole genome sequences of known SC2r-CoVs and three SC1r-CoVs (SARS-CoV-1 Tor2 (NC_004718.3), WIV1 (KF367457.1), and LYRa11 (KF569996.1) were downloaded from NCBI GenBank (https://www.ncbi.nlm.nih.gov/; download date, March 24, 2023). Also, the nucleotide sequences of ACE2 of various species were collected from NCBI GenBank (download date, March 24, 2023). Information on the coronaviruses and ACE2 sequences are summarized in Table S1.

### Molecular phylogenetic analysis

ML tree of RBDs of SC2r-CoVs was constructed by the following procedures: Multiple sequence alignment (MSA) was constructed using MAFFT v7.511 (ref.^37^) with the default option. Alignment sites with <30% site coverage were excluded using trimAl v1.2rev59 (ref.^38^). Alignment sites corresponding to the RBD of SARS-CoV-2 S (e.g., nucleotide positions 22517–23185 in the SARS-CoV-2 Wuhan-Hu-1 strain) were used for the tree construction. The ML tree was constructed using RAxML-NG v1.1.0 (ref.^39^) under the GTR+G+I nucleotide substitution model with 100 bootstrap analyses.

ML tree of nucleotide sequences of Rhinolophus ACE2 was constructed by the following procedures: MSA was constructed using MUSCLE^40^ implemented in MEGA11 v.11.0.13 (ref.^41^). ML tree was constructed using ML method implemented in MEGA11 under the T92+G substitution model with 100 bootstrap analyses.

### Cell culture

HEK293 cells (a human embryonic kidney cell line; ATCC, CRL-1573), HEK293T cells (a human embryonic kidney cell line; ATCC, CRL-3216) and HOS-ACE2/TMPRSS2 cells, HOS cells (a human osteosarcoma cell line; ATCC CRL-1543) stably expressing human ACE2 and TMPRSS2 (ref.^42, 43^) were maintained in Dulbecco’s modified Eagle’s medium (high glucose) (Sigma-Aldrich, Cat# 6429-500ML) containing 10% fetal bovine serum and 1% penicillin-streptomycin (Sigma-Aldrich, Cat# P4333-100ML). HOS-ACE2/TMPRSS2 cells, HOS cells (a human osteosarcoma cell line; ATCC CRL-1543) stably expressing human ACE2 and TMPRSS2 (ref.^42, 43^). The HOS-TMPRSS2 cells that stably express Rhinolophus bat ACE2 were generated as described below (see “Generation of HOS-TMPRSS2 cells stably expressing a variety of ACE2 proteins” section) and maintained in Dulbecco’s modified Eagle’s medium (high glucose) (Sigma-Aldrich, Cat# 6429-500ML) containing 10% fetal bovine serum and 1% penicillin-streptomycin (Sigma-Aldrich, Cat# P4333-100ML).

### Plasmid construction

Plasmids expressing the codon-optimized S proteins of B236 (GenBank accession no. MZ937003.2) and B52 (GenBank accession no. MZ937000.1) were synthesized by a gene synthesis service (Fasmac). Plasmids expressing the ACE2 proteins of *R. sinicus* (GenBank accession no. KC881004.1), *R. ferrumequinum* (GenBank accession no. AB297479.1), *R. shameli* (GenBank accession no. MZ851782.1), *R. pearsonii* (GenBank accession no. EF569964.1), *R.pusillus* (GenBank accession no. GQ999938.1), *R. macrotis* (GenBank accession no. GQ999932.1), *R. affinis* (GenBank accession no. MT394225.1), hamster (GenBank accession no. XM_005074209.3), and pangolin (GenBank accession no. XM_017650263.2) were also synthesized by a gene synthesis service (Fasmac). For *R. cornutus* ACE2 (GenBank accession no. LC564973.1), pCAGGS-blast-RcACE2^44^ was provided. Plasmids expressing the derivatives of codon-optimized S proteins of B236 and B52 and Rhinolophus bat ACE2 were generated by site-directed overlap extension PCR using the primers listed in **Table S2**. The resulting PCR fragment was cloned into the KpnI-NotI site of backbone pCAGGS vector^45^ (for the S expression plasmids) or the BamHI/MluI site of pWPI-ACE2-zeo (for ACE2 expression plasmids)^43^ with 3×FLAG-tag at the C-terminus using In-Fusion® HD Cloning Kit (Takara, Cat# Z9650N). Nucleotide sequences were determined by DNA sequencing services (Eurofins), and the sequence data were analyzed by Sequencher v5.1 software (Gene Codes Corporation).

### Generation of HOS-TMPRSS2 cells stably expressing a variety of ACE2 proteins

To prepare lentiviral vectors expressing ACE2, HEK293T cells (2,000,000 cells) were cotransfected with 12 μg of pCAG-HIVgp, 10 μg of pCMV-VSV-G-RSV-Rev, and 17 μg of either pWPI-ACE2-zeo by the calcium phosphate method. After 12 hours of transfection, the culture medium was changed to fresh medium. After 48 hours of transfection, the culture supernatant including lentivector particles was collected. HOS-TMPRSS2 cells (100,000 cells) were then transduced with the ACE2-expressing lentiviral vector. After 48 hours post transduction, transduced cells were maintained for zeocin (50 μg/mL; Invivogen, Cat#ant-zn-1) selections for 14 days.

### Pseudovirus assay

Pseudovirus assay was performed as previously described^28–31, 43, 46–54^. Briefly, HIV-1-based, luciferase-expressing reporter viruses were pseudotyped with the S proteins of B236, B52 and their derivatives. HEK293T cells (3,000,000 cells) were cotransfected with 4 μg psPAX2-IN/HiBiT^55^, 4 μg pWPI-Luc2 (ref.^55^), and 2 μg plasmids expressing parental S or its derivatives using PEI Max (Polysciences, Cat# 24765-1) according to the manufacturer’s protocol. Two days posttransfection, the culture supernatants were harvested, and the pseudoviruses were stored at –80°C until use. For pseudovirus infection, the amount of input virus was normalized to the HiBiT value measured by Nano Glo HiBiT lytic detection system (Promega, Cat# N3040)], which indicates the amount of p24 HIV-1 antigen. For target cells, the HOS-TMPRSS2 cells stably expressing a variety of Rhinolophus bat ACE2 (**Fig. 1D**) and the HEK293 cells transfected with the plasmids expressing *R. pusillus* and *R. macrotis* ACE2 and their derivatives with TransIT-LT1 (Takara, Cat# MIR2300) (**Fig. 3B** and **3D**) were used. Two days postinfection, the infected cells were lysed with a Bright-Glo Luciferase Assay System (Promega, cat# E2620) and the luminescent signal was measured using a GloMax Explorer Multimode Microplate Reader (Promega).

### Association analysis between the B236 and B52 infection tropism and polymorphic sites in ACE2 among animals

The results of the B236 and B52 pseudoviral infection assay in the cells expressing ACE2 from Rhinolophus bats and other representative species were used. Of these, results for *R. affinis, R. cornutus, R. ferrumequinum, R. macrotis, R. pusillus, R. shameli, R. sinicus*, hamster, and pangolin were used: We excluded the data for human and *R. pearsonii* from the analysis because the data for these species deviated from the trend in which ACE2 proteins more sensitive to B236 infection are less sensitive to B52. First, we calculated relative infectivity (log_10_(B236 infectivity / B52 infectivity)) for each ACE2 protein. This elative infectivity score was used as an objective variable for the association analysis. We constructed the MSA of ACE2 amino acid sequences using MAFFT v7.511 (ref.^37^) with the default option. We used amino acid residues in each alignment site of the MSA as qualitative explanatory variables. Statistical significance of the association between the relative infectivity and amino acid residues in an ACE2 polymorphic site was evaluated using one-way ANOVA. The analysis was performed in R v4.2.1.

### Protein structure model

All protein structural analyses were performed using Discovery Studio 2021 (Dassault Systèmes BIOVIA). In **Fig. 2D** and **2E**, the crystal costructure of B236 S RBD and human ACE2 (PDB: 7PKI)^6^ was used as the template, and 40 homology models of the B52 S RBD were generated using Build homology model protocol MODELLER v9.24 (ref.^56^). Evaluation of the homology models were performed using PDF total scores and DOPE scores and the best model for the B52 S was selected. Homology models of ACE2 of *R. cornutus*, *R. pusillus*, *R. macrotis* or *R. sinicus* were generated with the same way as B52 S RBD. The crystal costructure of B236 S RBD and human ACE2 (PDB: 7PKI)^6^ were used. To predict interaction between S RBD and ACE2, the structure of human ACE2 was replaced by the homology models of ACE2 of *R. cornutus*, *R. pusillus*, *R. macrotis* or *R. sinicus*. In **Fig. 2F**, the structure of B236 S RBD was replaced by the homology model of B52 S RBD.

### Western blot

Western blot was performed as previously described^29, 30, 49, 51, 53, 57^. For the blot, the supernatants of HEK293T cells cotransfected with the S expression plasmids and HIV-1-based pseudovirus producing plasmids, or HEK293 cells cotransfected with bat ACE2 expression plasmids (see “Pseudovirus assay” section) were used. The harvested cells were washed and lysed in RIPA buffer (50□mM Tris-HCl buffer [pH 7.6], 150□mM NaCl, 1% Nonidet P-40, 0.5% sodium deoxycholate, 0.1% SDS), protease inhibitor cocktail (Nacalai Tesque, Cat# 03969-21)]. The lysates were diluted with 2 × sample buffer [100 mM Tris-HCl (pH 6.8), 4% SDS, 12% β-mercaptoethanol, 20% glycerol, 0.05% bromophenol blue]. Both samples were boiled for 10 m. Then, 10 μl samples were subjected to Western blot. In addition, 900 μl culture medium containing the pseudoviruses at 250 ng HIV-1 p24 antigen was layered onto 500 μl 20% sucrose in PBS and centrifuged at 20,000 g for 2 hours at 4°C. Pelleted virions were resuspended in 1 × sample buffer [50 mM Tris-HCl (pH 6.8), 2% SDS, 6% β-mercaptoethanol, 10% glycerol, 0.0025% bromophenol blue] and boiled for 10 m. For protein detection, the following antibodies were used: mouse anti-SARS-CoV-2 S monoclonal antibody (clone 1A9, GeneTex, Cat# GTX632604, 1:10,000), mouse anti-HIV-1 p24 monoclonal antibody (183-H12-5C, obtained from the HIV Reagent Program, NIH, Cat# ARP-3537, 1:1,000), mouse anti-alpha-tubulin (TUBA) monoclonal antibody (clone DM1A, Sigma-Aldrich, Cat# T9026, 1:10,000), horseradish peroxidase (HRP)-conjugated mouse anti-FLAG monoclonal antibody (clone M2, Sigma-Aldrich, Cat# A8592, 1:1,000), and HRP-conjugated horse anti-mouse IgG antibody (Cell Signaling, Cat# 7076S, 1:2,000). Chemiluminescence was detected using SuperSignal West Femto Maximum Sensitivity Substrate (Thermo Fisher Scientific, Cat# 34095), or Western Lightning Plus-ECL (PerkinElmer, Cat# NEL104001EA) according to the manufacturer’s instruction. Bands were visualized using ChemiDoc Touch Imaging System (Bio-Rad).

## Code availability

Computational codes used in the present study is available on the GitHub repository (https://github.com/TheSatoLab/BANAL_tropism).

## References

1. Zhou, P., Yang, X.L., Wang, X.G., et al. (2020). A pneumonia outbreak associated with a new coronavirus of probable bat origin. Nature 579, 270–273, 10.1038/s41586-020-2012-7.

2. Zhou, H., Chen, X., Hu, T., et al. (2020). A Novel Bat Coronavirus Closely Related to SARS-CoV-2 Contains Natural Insertions at the S1/S2 Cleavage Site of the Spike Protein. Curr Biol 30, 2196–2203 e2193, 10.1016/j.cub.2020.05.023.

3. Wacharapluesadee, S., Tan, C.W., Maneeorn, P., et al. (2021). Evidence for SARS-CoV-2 related coronaviruses circulating in bats and pangolins in Southeast Asia. Nat Commun 12, 972, 10.1038/s41467-021-21240-1.

4. Zhou, H., Ji, J., Chen, X., et al. (2021). Identification of novel bat coronaviruses sheds light on the evolutionary origins of SARS-CoV-2 and related viruses. Cell 184, 4380–4391 e4314, 10.1016/j.cell.2021.06.008.

5. Delaune, D., Hul, V., Karlsson, E.A., et al. (2021). A novel SARS-CoV-2 related coronavirus in bats from Cambodia. Nat Commun 12, 6563, 10.1038/s41467-021-26809-4.

6. Temmam, S., Vongphayloth, K., Baquero, E., et al. (2022). Bat coronaviruses related to SARS-CoV-2 and infectious for human cells. Nature 604, 330–336, 10.1038/s41586-022-04532-4.

7. Murakami, S., Kitamura, T., Suzuki, J., et al. (2020). Detection and Characterization of Bat Sarbecovirus Phylogenetically Related to SARS-CoV-2, Japan. Emerg Infect Dis 26, 3025–3029, 10.3201/eid2612.203386.

8. Xiao, K., Zhai, J., Feng, Y., et al. (2020). Isolation of SARS-CoV-2-related coronavirus from Malayan pangolins. Nature 583, 286–289, 10.1038/s41586-020-2313-x.

9. Lam, T.T., Jia, N., Zhang, Y.W., et al. (2020). Identifying SARS-CoV-2-related coronaviruses in Malayan pangolins. Nature 583, 282–285, 10.1038/s41586-020-2169-0.

10. Liu, K., Pan, X., Li, L., et al. (2021). Binding and molecular basis of the bat coronavirus RaTG13 virus to ACE2 in humans and other species. Cell 184, 3438–3451 e3410, 10.1016/j.cell.2021.05.031.

11. Niu, S., Wang, J., Bai, B., et al. (2021). Molecular basis of cross-species ACE2 interactions with SARS-CoV-2-like viruses of pangolin origin. EMBO J 40, e107786, 10.15252/embj.2021107786.

12. Wrobel, A.G., Benton, D.J., Xu, P., et al. (2021). Structure and binding properties of Pangolin-CoV spike glycoprotein inform the evolution of SARS-CoV-2. Nat Commun 12, 837, 10.1038/s41467-021-21006-9.

13. Li, P., Hu, J., Liu, Y., et al. (2023). Effect of polymorphism in Rhinolophus affinis ACE2 on entry of SARS-CoV-2 related bat coronaviruses. PLoS Pathog 19, e1011116, 10.1371/journal.ppat.1011116.

14. Yan, H., Jiao, H., Liu, Q., et al. (2021). ACE2 receptor usage reveals variation in susceptibility to SARS-CoV and SARS-CoV-2 infection among bat species. Nat Ecol Evol 5, 600–608, 10.1038/s41559-021-01407-1.

15. Guo, H., Hu, B.J., Yang, X.L., et al. (2020). Evolutionary Arms Race between Virus and Host Drives Genetic Diversity in Bat Severe Acute Respiratory Syndrome-Related Coronavirus Spike Genes. J Virol 94, 10.1128/JVI.00902-20.

16. Shang, J., Ye, G., Shi, K., et al. (2020). Structural basis of receptor recognition by SARS-CoV-2. Nature 581, 221–224, 10.1038/s41586-020-2179-y.

17. Maginnis, M.S. (2018). Virus-Receptor Interactions: The Key to Cellular Invasion. J Mol Biol 430, 2590–2611, 10.1016/j.jmb.2018.06.024.

18. Shi, Y., Wu, Y., Zhang, W., et al. (2014). Enabling the ‘host jump’: structural determinants of receptor-binding specificity in influenza A viruses. Nat Rev Microbiol 12, 822–831, 10.1038/nrmicro3362.

19. de Graaf, M., and Fouchier, R.A. (2014). Role of receptor binding specificity in influenza A virus transmission and pathogenesis. EMBO J 33, 823–841, 10.1002/embj.201387442.

20. Russell, R.M., Bibollet-Ruche, F., Liu, W., et al. (2021). CD4 receptor diversity represents an ancient protection mechanism against primate lentiviruses. Proc Natl Acad Sci U S A 118, 10.1073/pnas.2025914118.

21. Gautam, R., Gaufin, T., Butler, I., et al. (2009). Simian immunodeficiency virus SIVrcm, a unique CCR2-tropic virus, selectively depletes memory CD4+ T cells in pigtailed macaques through expanded coreceptor usage in vivo. J Virol 83, 7894–7908, 10.1128/JVI.00444-09.

22. Chen, Z., Kwon, D., Jin, Z., et al. (1998). Natural infection of a homozygous delta24 CCR5 red-capped mangabey with an R2b-tropic simian immunodeficiency virus. J Exp Med 188, 2057–2065, 10.1084/jem.188.11.2057.

23. Wetzel, K.S., Yi, Y., Elliott, S.T.C., et al. (2017). CXCR6-Mediated Simian Immunodeficiency Virus SIVagmSab Entry into Sabaeus African Green Monkey Lymphocytes Implicates Widespread Use of Non-CCR5 Pathways in Natural Host Infections. J Virol 91, 10.1128/JVI.01626-16.

24. Riddick, N.E., Wu, F., Matsuda, K., et al. (2015). Simian Immunodeficiency Virus SIVagm Efficiently Utilizes Non-CCR5 Entry Pathways in African Green Monkey Lymphocytes: Potential Role for GPR15 and CXCR6 as Viral Coreceptors. J Virol 90, 2316–2331, 10.1128/JVI.02529-15.

25. Elliott, S.T., Wetzel, K.S., Francella, N., et al. (2015). Dualtropic CXCR6/CCR5 Simian Immunodeficiency Virus (SIV) Infection of Sooty Mangabey Primary Lymphocytes: Distinct Coreceptor Use in Natural versus Pathogenic Hosts of SIV. J Virol 89, 9252–9261, 10.1128/JVI.01236-15.

26. Ng, M., Ndungo, E., Kaczmarek, M.E., et al. (2015). Filovirus receptor NPC1 contributes to species-specific patterns of ebolavirus susceptibility in bats. Elife 4, 10.7554/eLife.11785.

27. Takadate, Y., Kondoh, T., Igarashi, M., et al. (2020). Niemann-Pick C1 Heterogeneity of Bat Cells Controls Filovirus Tropism. Cell Rep 30, 308–319 e305, 10.1016/j.celrep.2019.12.042.

28. Kimura, I., Kosugi, Y., Wu, J., et al. (2022). The SARS-CoV-2 Lambda variant exhibits enhanced infectivity and immune resistance. Cell Rep 38, 110218, 10.1016/j.celrep.2021.110218.

29. Yamasoba, D., Kimura, I., Nasser, H., et al. (2022). Virological characteristics of the SARS-CoV-2 Omicron BA.2 spike. Cell 185, 2103-2115.e2119, 10.1016/j.cell.2022.04.035.

30. Kimura, I., Yamasoba, D., Tamura, T., et al. (2022). Virological characteristics of the novel SARS-CoV-2 Omicron variants including BA.4 and BA.5. Cell 185, 3992-4007.e3916.

31. Saito, A., Tamura, T., Zahradnik, J., et al. (2022). Virological characteristics of the SARS-CoV-2 Omicron BA.2.75 variant. Cell Host Microbe 30, 1540–1555.e1515, 10.1016/j.chom.2022.10.003.

32. Ito, J., Suzuki, R., Uriu, K., et al. (2022). Convergent evolution of the SARS-CoV-2 Omicron subvariants leading to the emergence of BQ.1.1 variant. BioRxiv doi: https://doi.org/10.1101/2022.1112.1105.519085.

26. Tamura, T., Ito, J., Uriu, K., et al. (2022). Virological characteristics of the SARS-CoV-2 XBB variant derived from recombination of two Omicron subvariants. BioRxiv doi: https://doi.org/10.1101/2022.1112.1127.521986.

34. Wang, Q., Guo, Y., Iketani, S., et al. (2022). Antibody evasion by SARS-CoV-2 Omicron subvariants BA.2.12.1, BA.4, & BA.5. Nature 10.1038/s41586-022-05053-w.

35. Gu, H., Chen, Q., Yang, G., et al. (2020). Adaptation of SARS-CoV-2 in BALB/c mice for testing vaccine efficacy. Science 369, 1603–1607, 10.1126/science.abc4730.

36. Kumar, S., Suleski, M., Craig, J.M., et al. (2022). TimeTree 5: An Expanded Resource for Species Divergence Times. Mol Biol Evol 39, 10.1093/molbev/msac174.

37. Katoh, K., and Standley, D.M. (2013). MAFFT multiple sequence alignment software version 7: improvements in performance and usability. Mol Biol Evol 30, 772–780, 10.1093/molbev/mst010.

38. Capella-Gutierrez, S., Silla-Martinez, J.M., and Gabaldon, T. (2009). trimAl: a tool for automated alignment trimming in large-scale phylogenetic analyses. Bioinformatics 25, 1972–1973, 10.1093/bioinformatics/btp348.

39. Kozlov, A.M., Darriba, D., Flouri, T., et al. (2019). RAxML-NG: a fast, scalable and user-friendly tool for maximum likelihood phylogenetic inference. Bioinformatics 35, 4453–4455, 10.1093/bioinformatics/btz305.

40. Edgar, R.C. (2004). MUSCLE: a multiple sequence alignment method with reduced time and space complexity. BMC Bioinformatics 5, 113, 10.1186/1471-2105-5-113.

41. Tamura, K., Stecher, G., and Kumar, S. (2021). MEGA11: Molecular Evolutionary Genetics Analysis Version 11. Mol Biol Evol 38, 3022–3027, 10.1093/molbev/msab120.

42. Ozono, S., Zhang, Y., Ode, H., et al. (2021). SARS-CoV-2 D614G spike mutation increases entry efficiency with enhanced ACE2-binding affinity. Nat Commun 12, 848, 10.1038/s41467-021-21118-2.

43. Ferreira, I., Kemp, S.A., Datir, R., et al. (2021). SARS-CoV-2 B.1.617 mutations L452R and E484Q are not synergistic for antibody evasion. J Infect Dis 224, 989-994, 10.1093/infdis/jiab368.

44. Murakami, S., Kitamura, T., Matsugo, H., et al. (2022). Isolation of Bat Sarbecoviruses, Japan. Emerg Infect Dis 28, 2500–2503, 10.3201/eid2812.220801.

45. Niwa, H., Yamamura, K., and Miyazaki, J. (1991). Efficient selection for high-expression transfectants with a novel eukaryotic vector. Gene 108, 193–199, 10.1016/0378-1119(91)90434-d.

46. Motozono, C., Toyoda, M., Zahradnik, J., et al. (2021). SARS-CoV-2 spike L452R variant evades cellular immunity and increases infectivity. Cell Host Microbe 29, 1124–1136, 10.1016/j.chom.2021.06.006.

47. Uriu, K., Kimura, I., Shirakawa, K., et al. (2021). Neutralization of the SARS-CoV-2 Mu variant by convalescent and vaccine serum. N Engl J Med 385, 2397–2399, 10.1056/NEJMc2114706.

48. Fujita, S., Kosugi, Y., Kimura, I., et al. (2022). Structural Insight into the Resistance of the SARS-CoV-2 Omicron BA.4 and BA.5 Variants to Cilgavimab. Viruses 14, 2677.

49. Suzuki, R., Yamasoba, D., Kimura, I., et al. (2022). Attenuated fusogenicity and pathogenicity of SARS-CoV-2 Omicron variant. Nature 603, 700–705, 10.1038/s41586-022-04462-1.

50. Uriu, K., Cardenas, P., Munoz, E., et al. (2022). Characterization of the immune resistance of SARS-CoV-2 Mu variant and the robust immunity induced by Mu infection. J Infect Dis 10.1093/infdis/jiac053.

51. Saito, A., Irie, T., Suzuki, R., et al. (2022). Enhanced fusogenicity and pathogenicity of SARS-CoV-2 Delta P681R mutation. Nature 602, 300–306, 10.1038/s41586-021-04266-9.

52. Yamasoba, D., Kosugi, Y., Kimura, I., et al. (2022). Neutralisation sensitivity of SARS-CoV-2 omicron subvariants to therapeutic monoclonal antibodies. Lancet Infect Dis 22, 942–943, 10.1016/S1473-3099(22)00365-6.

53. Kimura, I., Yamasoba, D., Nasser, H., et al. (2022). The SARS-CoV-2 spike S375F mutation characterizes the Omicron BA.1 variant. iScience 25, 105720, 10.1016/j.isci.2022.105720.

54. Uriu, K., Ito, J., Zahradnik, J., et al. (2023). Enhanced transmissibility, infectivity, and immune resistance of the SARS-CoV-2 omicron XBB.1.5 variant. Lancet Infect Dis 23, 280-281, 10.1016/S1473-3099(23)00051-8.

55. Ozono, S., Zhang, Y., Tobiume, M., et al. (2020). Super-rapid quantitation of the production of HIV-1 harboring a luminescent peptide tag. J Biol Chem 295, 13023–13030, 10.1074/jbc.RA120.013887.

56. Fiser, A., Do, R.K., and Sali, A. (2000). Modeling of loops in protein structures. Protein Sci 9, 1753–1773, 10.1110/ps.9.9.1753.

57. Nasser, H., Shimizu, R., Ito, J., et al. (2022). Monitoring fusion kinetics of viral and target cell membranes in living cells using a SARS-CoV-2 spike-protein-mediated membrane fusion assay. STAR Protoc 3, 101773, 10.1016/j.xpro.2022.101773.

