## Supplementary figures and images for "Determination of the factors responsible for host tropism of SARS-CoV-2-related bat coronaviruses"

### Fig. S1

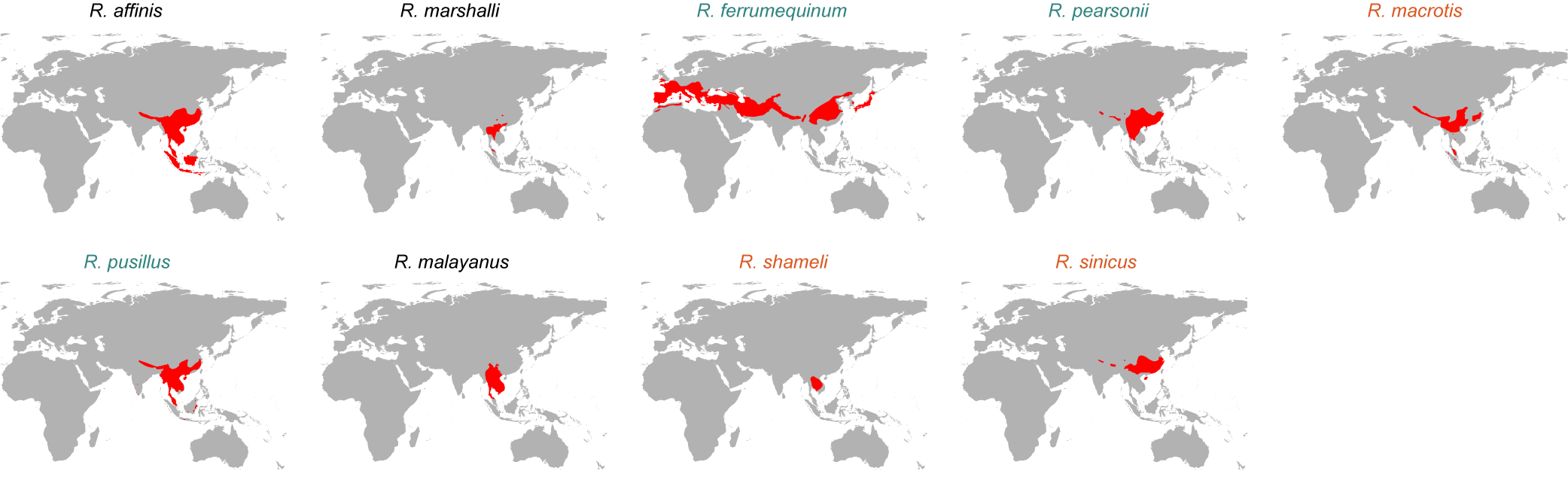

### Fig. S2

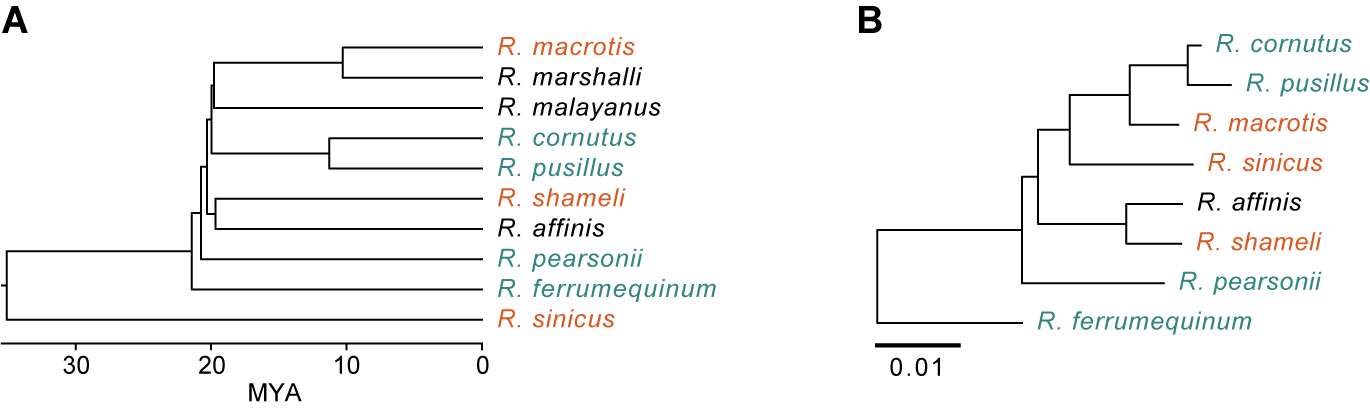

### Fig. S3

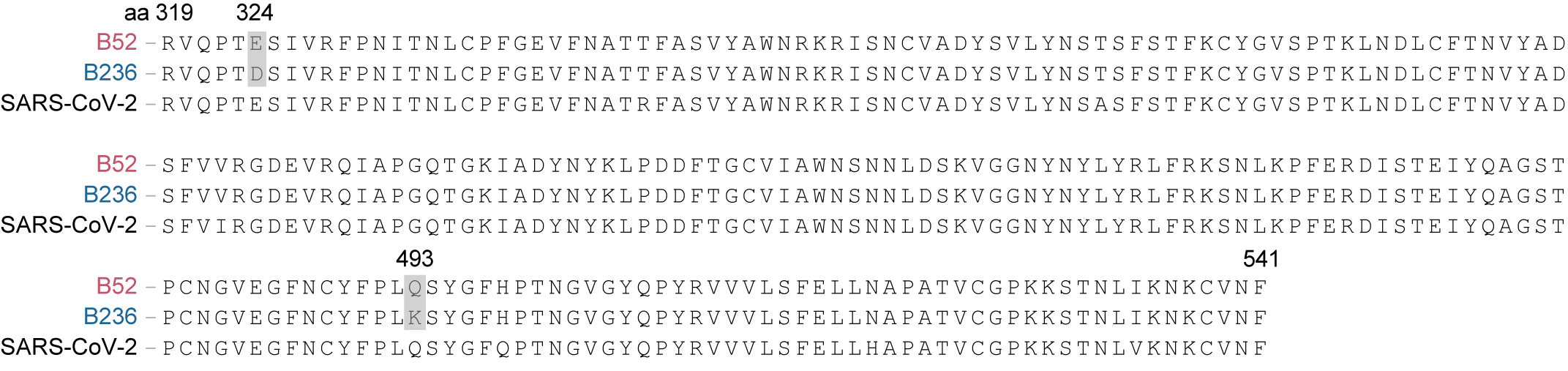
